# Linear contraction of stress fibers generates cell body rotation

**DOI:** 10.1101/2023.01.03.522661

**Authors:** Chika Okimura, Syu Akiyama, Yukinori Nishigami, Ryota Zaitsu, Tatsunari Sakurai, Yoshiaki Iwadate

**Affiliations:** Department of Biology, Yamaguchi University, Yamaguchi 753-8512, Japan; Research Institute for Electronic Science, Hokkaido University, Sapporo 001-0020, Japan; Graduate School of Life Science, Hokkaido University, Sapporo 001-0020, Japan; Department of Mathematical Engineering, Musashino University, Tokyo 135-8181, Japan

## Abstract

Wounds are healed by crawling migration of the epidermal cells around the injured area. Fish epidermal keratocytes that rapidly repair wounds comprise a frontal crescent-shaped lamellipodium and a rear rugby ball-shaped cell body. The cell body rotates like a wheel during migration. Stress fibers, which are bundles of contractile actomyosin filaments, are arranged along the seams of the rugby ball. Here we show the linear contraction of stress fibers to be the driving force for rotation. We constructed a mechanical model of the cell body that consisted of a soft cylinder with a contractile coil. From the motion of the model, it was predicted that contraction of the stress fibers would deform the soft cell body, as a result of which the deformed cell body would push against the substrate to generate torque. This prediction was confirmed by the observation of stress fiber dynamics in migrating cells. Linear-to-rotation conversion in migrating keratocytes is realized by simple soft-body mechanics. Conversion from linear motion to rotation is widely used in machines with moving parts, but requires somewhat complicated mechanics. An understanding of linear-to-rotation conversion in keratocytes has potential for use in the design of biomimetic soft robots.

## Introduction

Throughout the organismal life cycle, crawling cell migration plays an essential role in a variety of biological phenomena, including development^1^, wound healing^2^, immune function^3^, and cancer metastasis^4^. Wounds are healed by migration of surrounding epidermal cells to the injured area. The wound healing rate of fish skin is known to reach 500 *μ*m/h^5^, about 50 times faster than in human skin^6^. The crawling migration of fish epidermal keratocytes^7–9^ plays the main role in fast wound repair. A single keratocyte shows fast migration while maintaining its characteristic shape that consists of a frontal crescent-shaped lamellipodium and a rear rugby ball-shaped cell body^10–12^. At the leading edge of the lamellipodium, actin polymerization pushes the front^13–15^, whereas in the cell body, stress fibers composed of actomyosin are positioned to connect the left and right focal adhesions^16–19^. Contraction of the stress fibers retracts the rear^14,16,20^. However, the direction of the contractile forces is almost perpendicular to the direction of migration^13,20–23^, suggesting that the stress fibers play another role in cell advance.

Several investigators^14,24^ have reported that the cell bodies rotate in migrating keratocytes. Anderson et al. (1996) showed that cell membrane and nucleus also rotate, based on their observations of fluorescent beads in the inner part of the membrane and autofluorescence of the nucleus^24^. We also recently showed that multiple stress fibers, arranged like the seams of a rugby ball around the nucleus, rotate^25^. Ablation of some of the stress fibers induced the collapse of the left-right balance of the migrating cell. Rotation of the cell body, including stress fibers, appears to play an important role, at least in steering the cell while migrating.

If rotation of the cell body contributes to cell migration in the manner of a wheel, how it rotates is an interesting question. Anderson et al. (1996) observed that the translocation of the cell body continued after the arrest of lamellipodial protrusions by cytochalasin B^24^. We also observed that the stress fibers continued to rotate even after cutting off the leading edge of a migrating keratocyte^25^. These facts indicate that the cell body rotates autonomously, rather than passively by pseudopod extension.

Stress fibers are molecular machines that contract linearly like myofibrils^26–28^. We speculate that their linear contraction drives the rotation of the cell body, and that their linear motion is converted to rotation by as yet unknown mechanics. The construction of mechanical models that reproduce certain behaviors is a powerful technique for elucidating the mechanics of certain biological phenomena. For example, Nonaka et al. (2005) demonstrated, using a mechanical model composed of a rotating tilted wire that mimics a cilium^29^, that the cause of the nodal flow that creates left-right asymmetry in mice is the tilt of the cilia, which create a stream of viscous fluid.

In this study, we first confirmed that the rotation of stress fibers is accompanied by the slower rotation of the membrane and nucleus, suggesting that the stress fibers cause the rotation. We also constructed a mechanical model that mimics a keratocyte cell body using a soft silicone gel and a contractile coil. From a motion analysis of our mechanical model, a torque generation principle was posited: that the cell body, when deformed by stress fiber contraction, would push against the substrate. This prediction was confirmed by live cell imaging. Keratocytes use the elasticity of their cell body to achieve the conversion of linear motion into rotation.

## Results

### Rotation of the nucleus and stress fibers in a migrating keratocyte

In the rugby ball-shaped cell body of a migrating keratocyte (Fig. 1A and Supplementary Video 1), multiple stress fibers are arranged along the seams of the rugby ball around the nucleus (Fig. 1B)^25^. We first confirmed the rotation of the nucleus using sequential three-dimensional (3D) recordings (Supplementary Video 2), as Anderson et al. (1996) reported. A time series of top and bottom optical sections of the nucleus was compiled from sequential 3D images (Fig. 1C-F). Fluorescent spots of GFP-histone H1A at the top or bottom surfaces of the nucleus moved forward (Fig. 1D and the blue column in G, Intact) or rearward (Fig. 1F and the red column in G, Intact), respectively, in the cell frame of reference.

**Figure 1.**
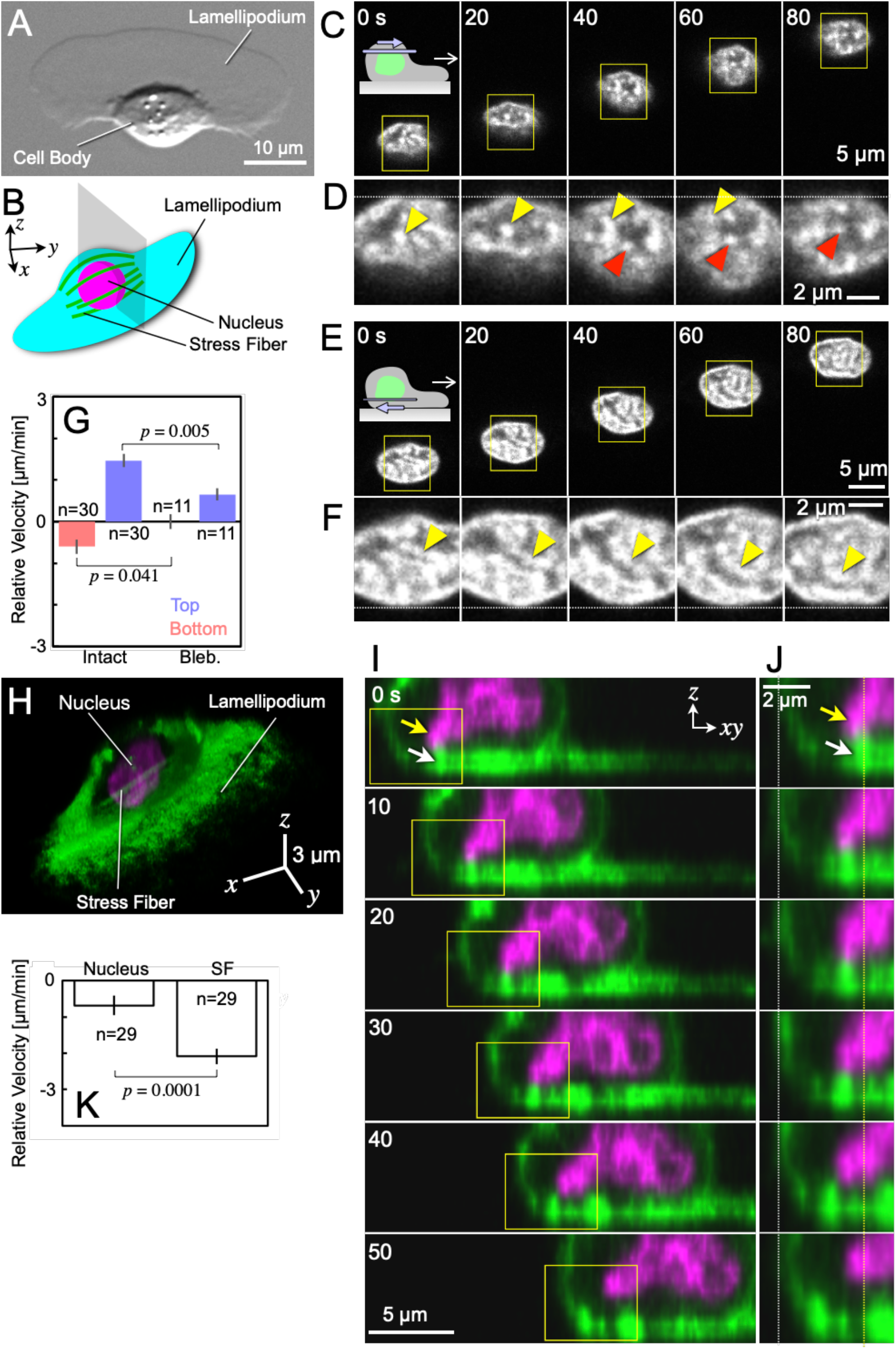
Rotation of the nucleus and stress fibers in a migrating keratocyte. (A) DIC image of a typical migrating keratocyte. (B) Schematic illustration of a migrating keratocyte. Stress fibers surround the nucleus. Gray: positions of the optical sections shown in I and J. (C - F) Sequential top (C and D) and bottom (E and F) images of a nucleus (GFP-Histone). D and F are enlarged images of the squares in C and E, respectively. The images are placed such that the anterior (D) and posterior (F) positions of the nucleus are respectively aligned at the dotted lines. The identical GFP-Histone fluorescent spots on the surface of the nucleus are indicated by the same-color arrowheads in each panel. The images in C -F are typical of 28 cells. (G) Relative velocities of fluorescent spots of GFP- or mCherry-Histone on the bottom and top surfaces of the nucleus in the cell frame of reference. Left two columns: untreated; right two columns: blebbistatin treatment. (H) A snapshot of a simultaneous 3D recording of the nucleus and stress fibers in a migrating untreated keratocyte. Histone (magenta, GFP-Histone) and F-actin (green, Alexa-phalloidin). (I) A nucleus and stress fibers at the grey optical sections in B, constructed from the simultaneous 3D recording as shown in H. White arrow: a stress fiber. Yellow arrow: the protrusion of a slightly distorted nucleus. (J) Enlarged images of the square in I. The images are placed such that the posterior edges of the cell are aligned at the white dotted line. Yellow dotted line: the position of the stress fiber at time 0 s. The images in I and J are typical of 29 cells. (K) Relative velocities of nucleus (Nucleus) and stress fibers (SF) in the cell frame of reference at the bottom surface of a migrating keratocyte. Error bars in G and K represent SEM. The *P* values were calculated using Student’s *t* test.

Next, to directly compare stress fiber rotation and nuclear rotation, we simultaneously recorded the dynamics of stress fibers and nucleus as sequential 3D images (Supplementary Video 3, top and Fig. 1H: snapshot from the images). The time series of perpendicular optical sections at the center of the cell (gray in Fig. 1B) were constructed from the sequential 3D images (Fig. 1I, J and Supplementary Video 3, bottom). At the bottom, a stress fiber (white arrows in Fig. 1I, J and Supplementary Video 3, bottom) moved rearward in the cell frame of reference. At the same time, a protrusion from a slightly distorted nucleus (yellow arrows in Fig. 1I, J and Supplementary Video 3, bottom) also moved rearward, dragged by the stress fibers (0 - 20 s in Fig. 1I and J). The protrusion then detached from the fiber (30 s in Fig.1I and J), and the protrusion stopped moving rearward and contracted (40 - 50 s in Fig. 1I and J). A statistical comparison of the velocities at the bottom of the cell (Fig. 1K) also shows that the rearward velocities of nuclei were lower than those of the stress fibers in the cell frame of reference, indicating that the rotation of the nucleus is not the cause of rotation of the stress fibers.

Treatment with low concentrations of blebbistatin, an inhibitor of myosin II ATPase, causes the disassembly of stress fibers but does not halt crawling cell migration^7,18,19,25,30^. We observed the dynamics of the nucleus in the blebbistatin-treated keratocytes during their migration. The velocities of fluorescent spots of mCherry-histone at both the top and the bottom surfaces significantly decreased in the cell frame of reference (Fig. 1G, Bleb.). These results suggest the rotation of stress fibers to be the driving force of the rotation of the nucleus. However, other possibilities cannot be entirely ruled out, since blebbistatin inhibits not only the polymerization of stress fibers but also other functions related to myosin II ATPase.

### Rotation of the cell membrane and stress fibers in a migrating keratocyte

Stress fibers are located inside the cell body and do not directly contact the extracellular substrate. If the rotation of the stress fibers contributes to cell migration, the cell membrane should also rotate in the same direction, at the same or at a slightly slower speed, in order to transfer the torque of stress fiber-rotation to the substrate. We stained the cell membrane of migrating keratocytes with Concanavalin A (Con A)-Alexa and recorded the dynamics of the bright spots at the top and bottom surface of the cell (Supplementary Video 4). The Con A spots at the top and bottom of the cell body moved forward and rearward in the cell frame of reference, respectively (Fig. 2A - D and E, Intact). Such movements, however, could not be detected in lamellipodia (Fig. 2B, Supplementary Video 4, Bottom). These results confirm the rotation of the membrane but not the lamellipodia, as indicated by Anderson et al. (1996)^24^.

**Figure 2.**
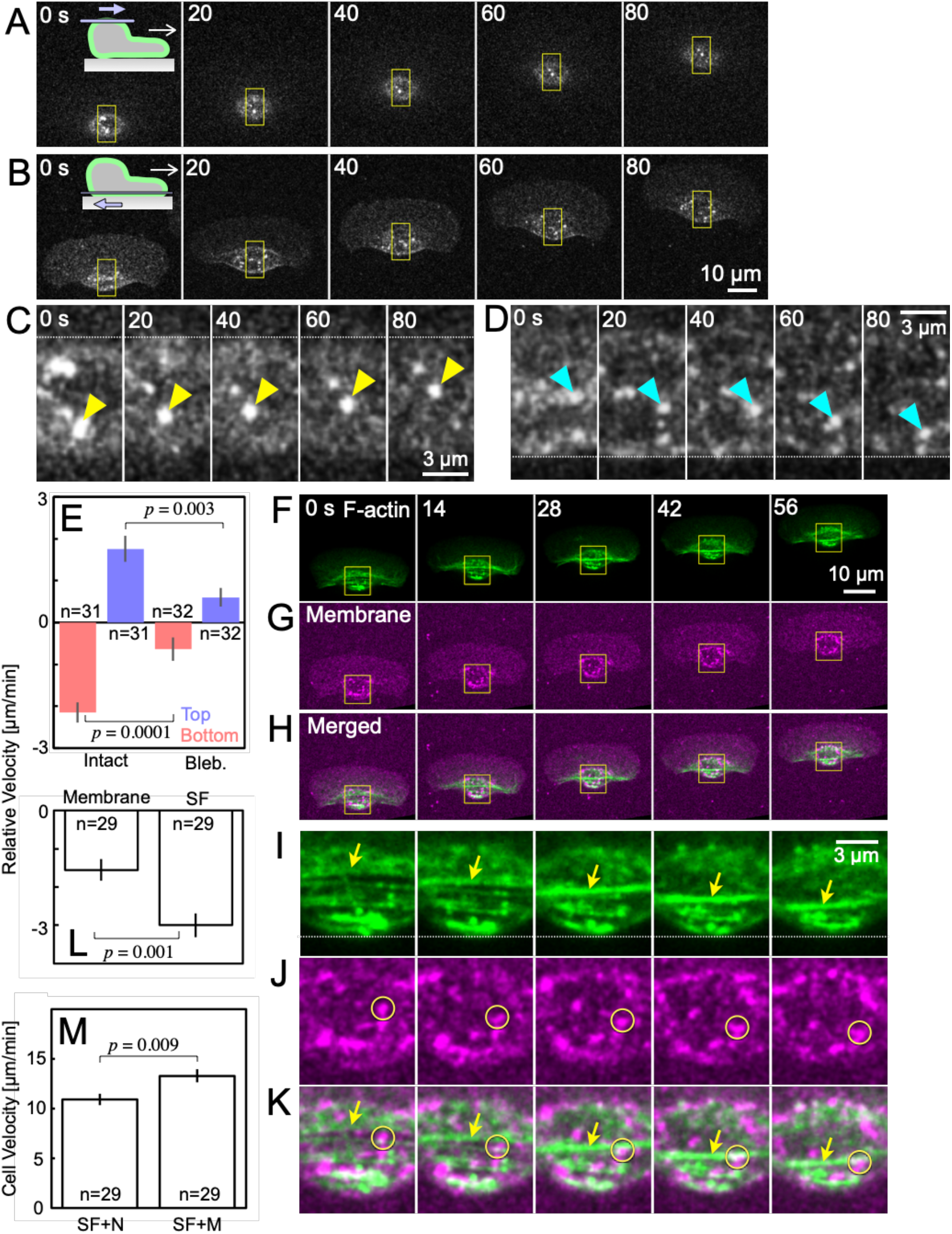
Rotation of cell membrane and stress fibers in a migrating keratocyte. (A and B) Sequential top (A) and bottom (B) membrane (Con A-Alexa) images of a migrating cell. (C and D) Enlarged images of the squares in A and B, respectively. The images are placed such that the anterior (C) and posterior (D) positions, respectively, of the cell body are aligned at the dotted lines. The identical Con A spots are indicated using the same-color arrowheads on each panel. The images in A- D are typical of 31 cells. (E) Relative velocities of the Con A spots at the bottom and top membrane in the cell frame of reference. Left two columns: untreated; right two columns: blebbistatin treatment. (F - H) Sequential images of F-actin (Alexa-phalloidin, F) and membrane (Con A-Alexa, G) at the bottom surface of a migrating keratocyte, and their merged images (H). (I - K) Enlarged images of the squares in F - H. The positions of the squares in the images at the same time of F - H are the same for all. The images of I - K are placed such that the posterior positions of the F-actin (I) are each aligned at the dotted lines. Yellow arrows in I and K indicate a stress fiber. The yellow circles in J and K enclose a single Con A spot. The images in F - K are typical of 20 cells. (L) Relative velocities of membrane (Membrane) and stress fibers (SF) in the cell frame of reference at the bottom surface of a migrating keratocyte. (M) Migration velocities of cells stained simultaneously for stress fibers and nucleus (SF+N, Fig. 1K) and those stained simultaneously for stress fibers and membrane (SF+M). Error bars in E, L and M represent SEM. The *P* values were calculated using Student’s *t* test.

To reveal whether rotation of the stress fibers causes membrane rotation or vice versa, we simultaneously recorded the dynamics of the F-actin and membrane at the bottom cell surface (Supplementary Video 5), and compared their velocities (Fig. 2F - L). During cell advance (Fig. 2F - H), a Con A spot (Fig. 2J and K, yellow circle) moved rearward at a somewhat slower speed than a stress fiber (Fig 2I and K, arrows) in the cell frame of reference. Statistically, the rearward velocity of Con A spots was slower than that of stress fibers (Fig. 2L), indicating that the membrane rotation is not the cause of stress fiber rotation. The rearward velocity of stress fibers is slower in Fig. 1K than in Fig. 2L. This is probably because after being extracted from a fish, keratocyte migration slows down over time^31^. Figure 2L shows cells one day after removal from the fish body, while Fig. 1K shows cells two days after extraction, kept to wait for the expression of GFP-histone. The migration velocity of the cells in Fig. 1K (Fig. 2M, SF+N) was slower than that in Fig. 2L (Fig. 2M, SF+M).

Next, we observed the Con A spots at the top and bottom of the cell body of migrating keratocytes whose stress fibers had been disassembled by treatment with low concentrations of blebbistatin. Blebbistatin treatment reduced the velocities of Con A spots at the top and bottom to 34% and 30%, respectively, of their original speed (Fig. 2E, Bleb.). These results suggest the rotation of stress fibers to be the driving force for membrane rotation, but possibly not the only driving force.

### Reproduction of the cell body rotation using a mechanical model

To test whether linear dynamics like stress fiber contraction can be converted into the rotation of the cell body, we constructed a mechanical model 1,500 × in scale (4.3 gf in weight, Fig. 3A and B): A shape memory alloy coil that mimics a stress fiber was embedded in the curved side of a soft cylindrical silicone gel, designed to model a cell body. The model was placed on a horizontal acrylic base plate. The adhesive force between the gel and the plate, measured as the force required to detach them, was 9.7 gf/cm^2^ (Supplementary Fig. 1A). Electric current was applied to the coil, whose resistance was 10 ± 0.3 Ω (SEM; n = 11) using a purpose-built electric power circuit (Supplementary Fig. 1B). In response to the application of an electric current of 140 mA (Fig. 3C), the coil contracted in response to Joule heating, causing the gel to deform. The model then rolled forward on its acrylic base plate (Fig. 3D, E and Supplementary Video 6).

**Figure 3.**
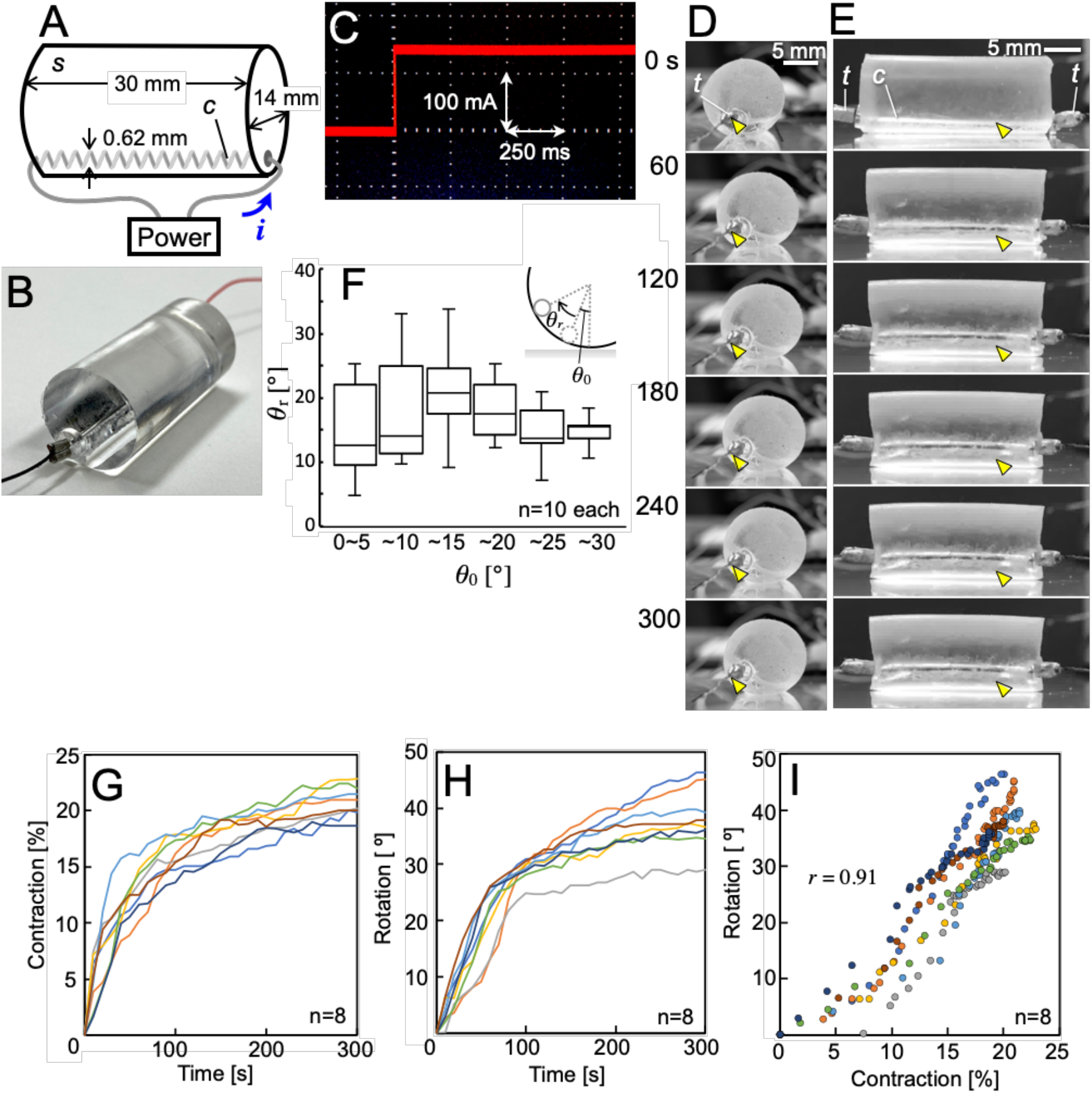
Rotation of keratocyte cell body model. (A) Schematic illustration of the model. It consists of a soft silicone gel cylinder (*s*) and a shape-memory alloy coil (*c*) that contracts when an electric current is applied. (B) Photo of the model. (C) Applied current (*i* in A) to the coil to cause its contraction. (D and E) Side view (D) and back view (E) of sequential images of the rotation. The yellow arrowheads in (D) and (E) indicate the coil position at 0 s and are located at the same position in each series of pictures in (D) and (E). Terminals (*t*) connecting the coil and the electric wire and a coil (*c*) are shown. (F) Relationship between the position of the coil contraction (*θ*_0_) and the angle of rotation (*θ*_r_). Inset: schematic illustration of *θ*_0_ and *θ*_r_. The top and bottom edges of the perpendicular line indicate the maximum and minimum values, the top and bottom of the square indicate the 75th and 25th percentiles, and the horizontal lines inside the squares show the median value. (G and H) Time courses of the coil contraction (G) and the angle of rotation (H) of the model. (I) Contraction-to-rotation curve made from (G) and (H).

The movement characteristics of this model were then investigated in detail. We first clarified the relationship between the position where the coil contacts the gel and the angle of rotation of the gel cylinder (*θ*_0_ and *θ*_r_ in Fig. 3F). The maximum angle of rotation was obtained by the coil contraction at a position of 10 - 15º from the vertical. Thus, in the following experiments, the coil was caused to contract at the 15º position. From the time-courses of the contraction ratio of the coil (Fig. 3G), which was defined as the linear distance between the two terminals of the contracted coil divided by the original distance, and the angle of rotation of the gel cylinder (Fig. 3H), a contraction-to-rotation relationship (Fig. 3I) was obtained. The correlation coefficient (*r*) between contraction and rotation was 0.91, indicating that contraction of the coil induces rotation of the gel

Vehicles must exert traction forces on the substrate to be able to move. The distribution of traction forces depends on the principle of motion of the given vehicle. Comparing the traction force maps exerted by each vehicle is thus a useful technique for comparing the principles of different vehicles. The traction force map exerted by our model (Fig. 4A and Supplementary Video 7) was almost same as that exerted by a migrating keratocyte (Fig. 4B and Supplementary Video 8). In both cases, traction forces are exerted mainly at the left and right edges toward the center. We constructed another model of a typical vehicle with front wheel drive and a trailing rear wheel (Fig. 4C). The wheels are made not of soft silicone but of hard acrylic, and their diameter and length are the same as our model of a keratocyte cell body (Fig. 3A and B). The traction force map of our model (Fig. 4A) was completely different from those of the vehicle model with front wheel drive and a trailing rear wheel (Fig. 4D, E and Supplementary Video 9). These results suggest that our mechanical model (Fig. 3A and B) accurately reproduces the rotation of keratocyte stress fibers and that both rotate according to the same mechanical principles.

**Figure 4.**
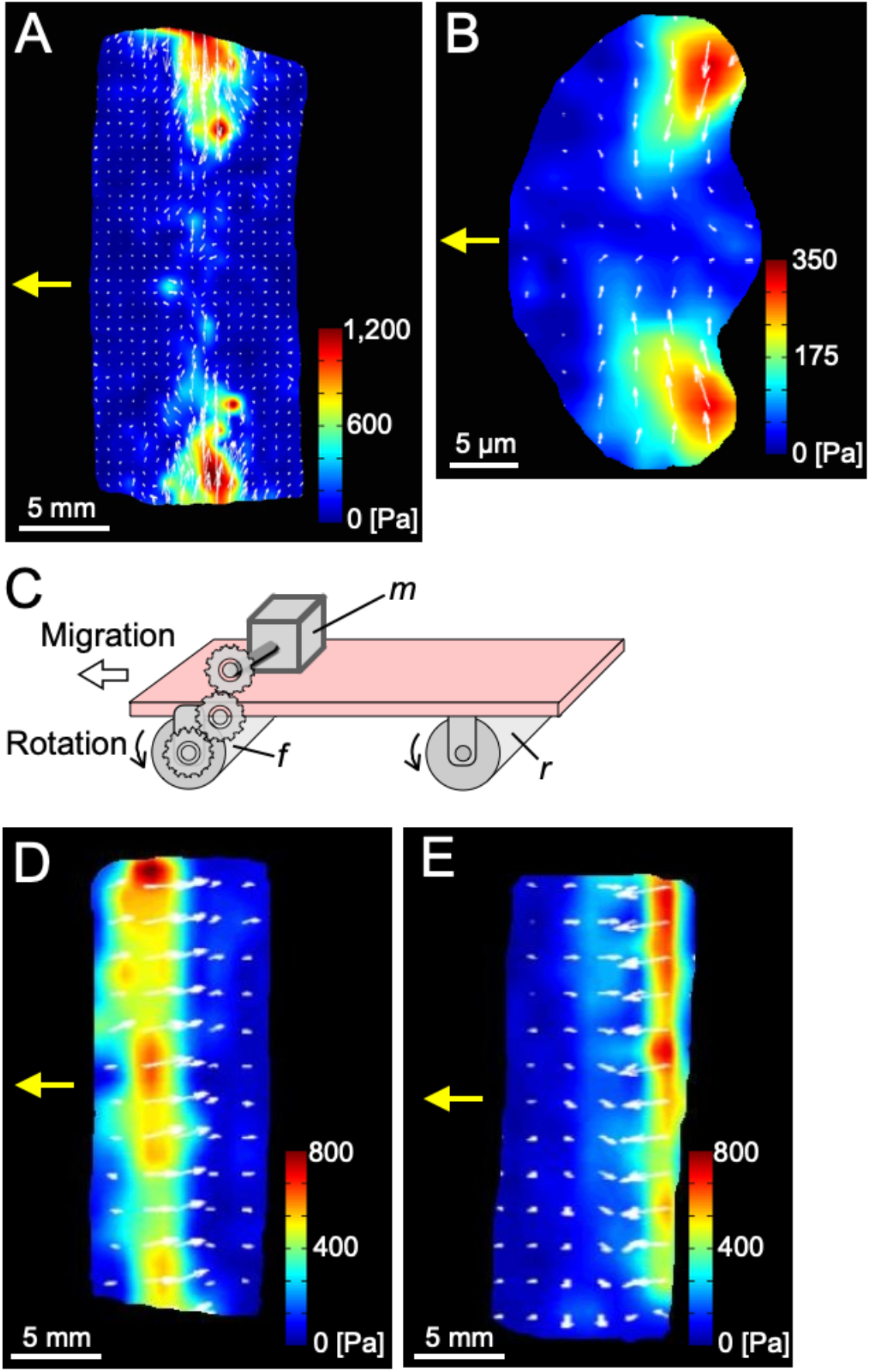
Traction force measurements. (A) Traction forces exerted by the mechanical model of the keratocyte cell body. The image in A is typical of 16 experiments. (B) Traction forces exerted by a migrating keratocyte. The image in B is typical of 17 cells. (C) Schematic illustration of a front-wheel-drive vehicle with a front wheel drive (*f*) with a motor (*m*) and a rear trailing wheel (*r*). (D and E) Traction forces exerted by the front (D) and the rear wheel (E) of the vehicle. The images in D and E are typical of 16 experiments. The direction and length of the white arrows in A, B, D and E respectively indicate the direction and relative magnitude of the forces there. Yellow arrows indicate the direction of migration of each model.

### Rotation of the model is induced by pushing against the substrate

To reveal how the model converts coil contraction into rotation of the gel, we accurately evaluated the deformation of the gel. By applying lime powder to the gel surface, the adhesive force of the gel was reduced to close to zero. Four glass rods were placed on the substrate parallel to the contraction direction of the coil without attaching them, and the model was placed on them such that *θ*_0_ was 90º (Supplementary Fig. 2A). In this situation, the contraction of the coil induces the deformation of the gel without being restricted by the adhesion of the gel to the substrate, but does not induce rotation of the gel. The images of the model before and after the coil contraction (Supplementary Fig. 2B and C) were taken from directly above (arrow in Supplementary Fig. 2A). From the images, the displacement of the centroid of the gel (*l* in Supplementary Fig. 2D) was calculated using the following formula.

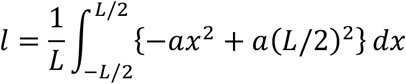

The centerline (blue line in Supplementary Fig. 2D) was approximated as a quadratic function. The term *a* is the constant of the function. The displacement of the centroid of the gel was proportional to the ratio of the coil contraction (Supplementary Fig. 2E). Since the maximum contraction ratio of the migrating mechanical model was about 20% (Fig. 3G), from the graph, we estimated the maximum displacement of the centroid to be 1.4 mm (Supplementary Fig. 2E).

From these results, we considered two possibilities as the cause of the model’s rotation; (1) displacement of the centroid and/or (2) pushing against the substrate caused by deformation of the gel. First, we tested the former possibility. From the 1.4-mm displacement of the centroid, it was calculated that when the substrate is tilted 2.5º (*ψ* in Supplementary Fig. 2F), the center of gravity of the new centroid position moves toward the grounding point of the gel. In this situation, the displacement of the centroid should not affect the rotation. We then observed whether or not the model would rotate on a substrate tilted by 2.5º (Fig. 5A). Coil contraction occurred normally (Fig. 5B). Rotation of the gel also occurred (Fig. 5C and D) as it did on the horizontal substrate (Fig. 3H and I). The correlation coefficient (*r*) between contraction and rotation was 0.95, indicating that displacement of the centroid does not contribute in any way to the rotation.

**Figure 5.**
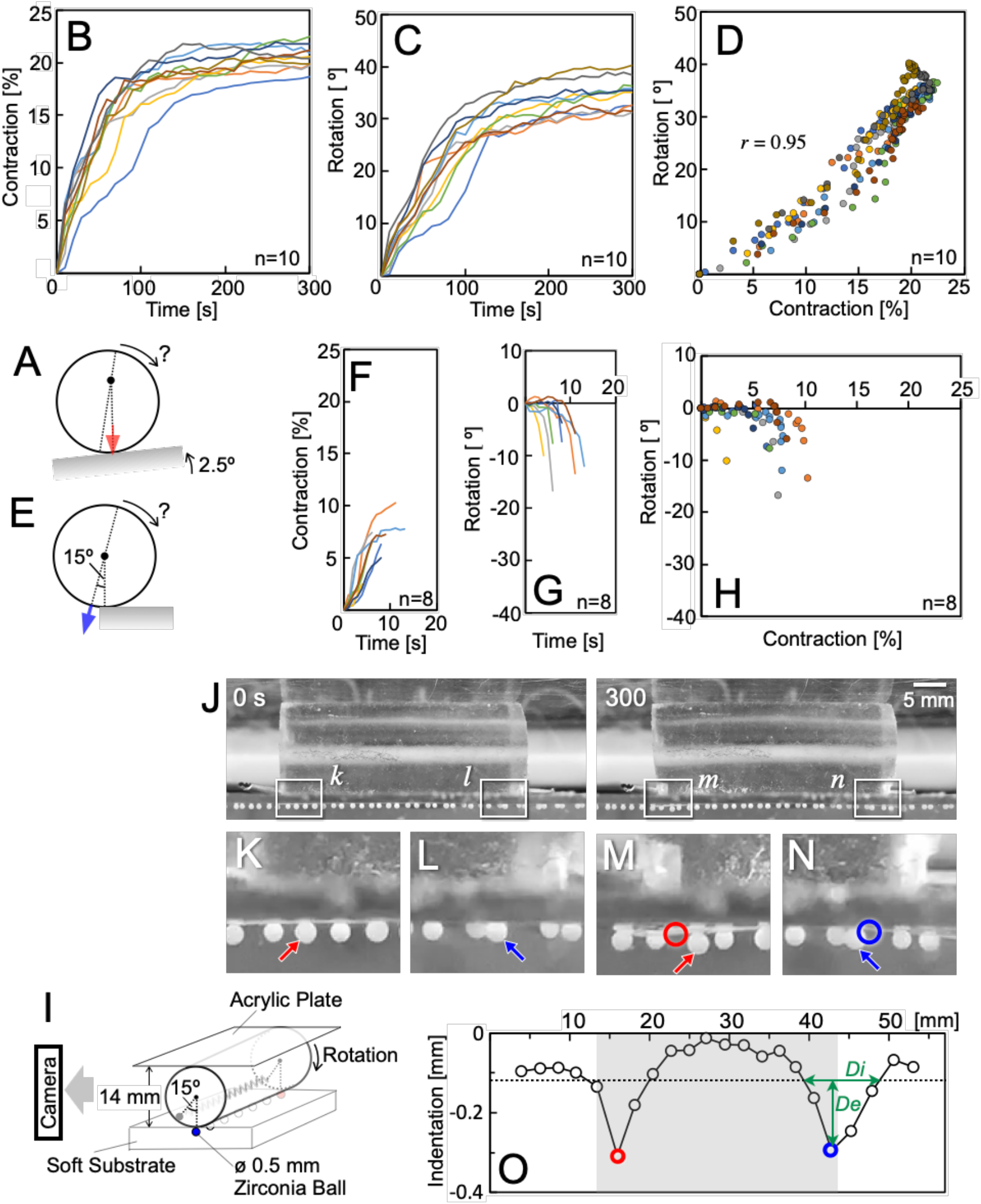
Evaluation of the mechanics of the rotation of the mechanical model. (A) Schematic illustration of the experiments on an uphill substrate. The tilt angle was set to 2.5º such that the gravitational pull on the displaced centroid (dot) was vertically downward (red arrow). (B and C) Time courses of the coil contraction (B) and the angle of rotation (C) of the model on the uphill substrate. (D) Contraction-to-rotation curve made from (B) and (C). (E) Schematic illustration of the experiments at the edge of a horizontal substrate. The gel is expected to exert a pushing force in the direction of the blue arrow. (F and G) Time courses of the coil contraction (F) and the angle of rotation (G) of the model at the edge of the substrate. (H) Contraction-to-rotation plots made from (F) and (G). (I) A schematic illustration of the experiment for detecting that the gel pushes the substrate. The height of the space in which the gel can move was limited to 14 mm (= diameter of the gel). Displacement of the zirconia balls embedded in the surface of the soft substrate was detected. (J) Mechanical model before (0 s) and after (300 s) contraction of the coil. (K - N) Enlarged images of *k* - *n* in J. Balls indicated by the arrows of the same color in K - N are the same. The circles in M and N show the same ball positions indicated by the same-color arrows in K and L. (O) Vertical components of ball displacements. Data for the balls indicated by arrows in K and L are shown as circles of the same color. The grey area shows the position of the model before contraction of the coil. Depth (*De*) and diameter (*Di*) of the indentation were defined based on the average height of substrate (dotted line). The images in J - N are typical of 15 experiments.

Then, to test the second possibility, we placed our model at the edge of the substrate (Fig. 5E). In this situation, the pushing force (blue arrow in Fig. 5E) cannot transmit to the substrate. The coil contraction began normally (Fig. 5F). However, the gel fell from the edge instead of rotating (Fig. 5G and H), suggesting that pushing against the substrate is the cause of the rotation. We then tested whether the gel would always push against the substrate. The mechanical model was placed on a soft elastic substrate that had a Young’s modulus of 5.1 kPa (Fig. 5I). The height of the space where the mechanical model could move was limited to 14 mm, identical to the diameter of the gel, using an acrylic plate. On contraction of the coil, the gel deformed and the substrate showed indentations just below both ends of the gel (Fig. 5J - O, Supplementary Fig. 3 and Supplementary Video 10). The depth and the radius of the indentations were 0.187 ± 0.0367 mm (n = 6) and 1.73 ± 113 mm (n = 6), respectively. From those values, the force pushing against the substrate is calculated to be 0.96 mN (= 98 mgf). These results indicate that our mechanical model generates the torque for its rotation from coil contraction by pushing against the substrate rather than via displacement of the centroid.

### Keratocyte cell bodies push against the substrate to effect rotation

The similarity of the traction force distribution between our mechanical model (Fig. 4A) and keratocytes (Fig. 4B) suggests that the rotation of the cell body in keratocytes is realized via the same mechanics as that of our model. It is essential for rotation for our model to push against the substrate as a result of its deformation (Fig. 5). We dispersed keratocytes on a soft substrate with a Young’s modulus of 0.1 kPa. The height of the space in which the cells could move was limited to the same height as the cell, by covering them with an agar sheet (Fig. 6A). Under these conditions, the stress fibers did not rotate (Fig. 6B and Supplementary Video 11), and the indentations in the substrate were detected at the left and right ends of the cell body just before the cell body passed by (Fig. 6C, red and yellow squares, D, Left and Right, and Supplementary Video 12). The depth of the indentations was 225 ± 0.0125 nm (n = 5). From that value, the force pushing against the substrate is calculated to be 32 ± 2.6 pN (n = 5). On the other hand, since the cells were covered with an agarose sheet, the substrate just below the middle part of the cell body showed indentations due to the volume effect the entire time the cell body was on the area (Fig. 6C, blue squares, D, Middle, and Supplementary Video 12). These results suggest that the cell body rotation of keratocyte requires pushing forces against the substrate at the both ends of the cell body, created by contraction of the stress fibers.

**Figure 6.**
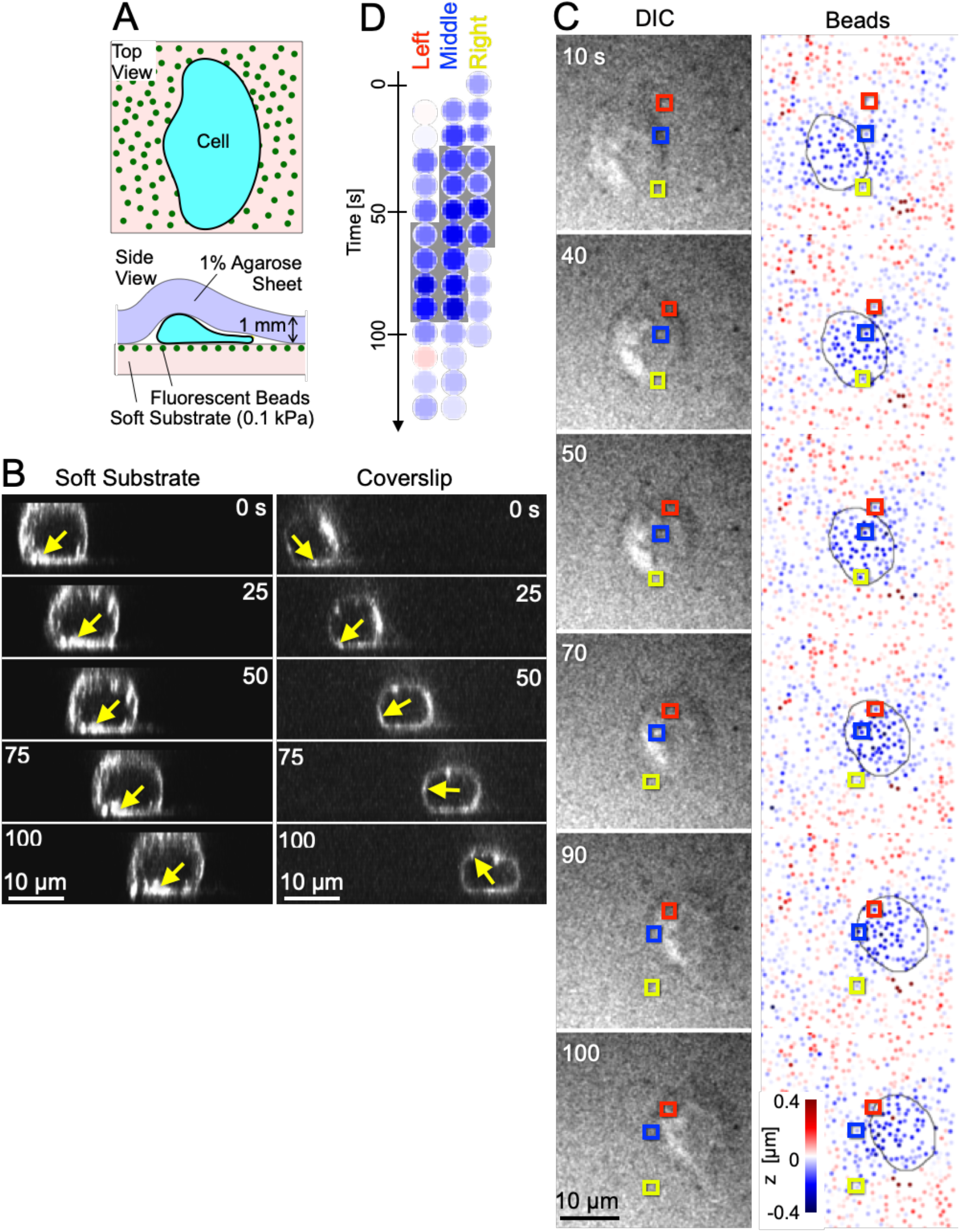
Migration of keratocytes on a soft substrate. (A) Schematic illustration of the experiments. (B) No stress fiber rotation. Longitudinal image sequence of F-actin made from a 3D recording of a migrating keratocyte on a soft substrate with Young’s modulus of 0.1 kPa (left, SPY555) and on a coverslip (right, Alexa-phalloidin). The position of the optical sections is the same as in Fig. 1I. Stress fibers do not detach from the substrate at the rear of the cell body (left, yellow arrows) or do (right, yellow arrows). The left and right images in B are typical of 17 and 13 cells, respectively. (C) Sequential DIC images of a migrating cell (left) and pseudocolor images of bead heights as the cell passes by (right). Three beads passing under the left, middle, and right of the cell are respectively indicated by red, blue and yellow squares. The images in C are typical of 9 events from 8 cells. (D) Enlarged sequential images of the three bead heights observed in C. Time periods when the beads are under the cell body are shown with a gray background. Images at times not in C are also displayed.

## Discussion

Numerous studies that combine mathematical modeling and biological experiments have revealed various biological phenomena^12,32–39^. These studies have a common and powerful strategy, i.e., mathematical model predictions that were confirmed by biological experimentation. We adopted the same approach in this study, using a mechanical model rather than a mathematical model. By combining mechanical modeling and biological experiments, we elucidated how stress fiber contractions are converted to cell body rotation (Supplementary Fig. 4). Our mechanical model predicted that the cell body would rotate by pushing against the substrate (Fig. 5 and Supplementary Video 10); then, as predicted, contraction of the stress fibers created indentations in the soft substrate just below both ends of the fibers (Fig. 6C, D and Supplementary Video 12), with no rotation of the stress fibers being observed (Fig. 6B and Supplementary Video 11). It is difficult to elucidate the mechanics by which the linear contraction of stress fibers is converted into rotational motion by means of biological experiments alone. Combining not only mathematical but also mechanical models with biological experiments is also useful.

The forces involved in cell migration such as leading-edge elongation are on the order of several nN^40^. The force of the cell body pushing against the substrate was only 32 pN. The force of the mechanical model pushing against the substrate (98 mgf) is also only 2.3% of its weight (4.3 gf), suggesting that strong forces are not necessary for this type of rotation.

Conversion from linear motion to rotation is used in many machines. For example, internal combustion engines that deliver rotational motion from the linear motion of pistons have a history of more than 100 years. The engine is made of rigid materials such as pistons, connecting rods, and a crankshaft. All of these complicated parts are needed to convert the linear motion of the pistons to the rotation of the crankshaft. Our mechanical model showed that linear contraction motion can be converted to rotational motion using only a contractile coil and a soft cylinder. This is a much simpler design than is used in existing machines. It appears that keratocytes have achieved a simple conversion strategy that uses their own flexibility, which is a characteristic of living organisms. Wounds are healed by the natural migration of surrounding epidermal cells to the injured area. Learning from this unique migration mechanism seen in fish keratocytes has the potential to lead to innovations in human wound healing therapy.

‘Soft robots’ that apply flexible features such as seen in living organisms to machines are a new concept that has attracted attention in recent years^41–44^. Our mechanical model not only elucidates the biological conversion mechanics from linear to rotational motion, but is also directly linked to biomimetics. In the near future, it may be possible to create a practical machine that converts contractile motion into rotational motion using simple mechanics consisting of multiple contractile fibers and a soft cylinder.

## Methods

### Cell culture

Primary cultures of keratocytes prepared from the scales of Central American cichlids (*Hypsophrys nicaraguensis*) as previously described^25^ were used throughout this study. Briefly, without sacrificing the fish, a few of their scales were removed and washed in culture medium (Leibovitz’s medium: L-15, L5520: Sigma-Aldrich supplemented with 10% fetal calf serum (Nichirei) and antibiotic/antimycotic solution (09366-44: Nacalai Tesque). The scales were placed external side up on the floor of a square chamber (18 × 18 mm and 2 mm in depth), the bottom of which was made of a 24 × 24 mm coverslip (No. 1, Matsunami). They were then covered with another small coverslip and allowed to adhere to the bottom coverslip for 1 h at 23 °C. Culture medium was then added to the chamber. After removal of the upper coverslip, the scales were kept at 23 °C again overnight to allow the keratocytes to spread from the scale. Cells were treated with 0.5 g/L trypsin and 0.53 mM EDTA (trypsin-EDTA, 32778-34: Nacalai Tesque) for 30 - 60 s to separate any cell-cell adhesions. All experiments were carried out in accordance with national guidelines and approved by Yamaguchi University Animal Use Committee.

### Transient transfection of keratocytes with eGFP- or mCherry-histone H1A construct

Polyethylene glycol (28254-85; Nacalai Tesque) was dissolved at 10 mM in Dulbecco’s phosphate-buffered saline with Ca^2+^ and Mg^2+^ (PBS++). This PEG medium was then added to the chamber, to the bottom of which the cells had adhered, immediately after removal of the culture medium (Leibovitz’s medium: L-15, 21083-027: Thermo Fisher) supplemented with 10% fetal calf serum (Nichirei) and antibiotic/antimycotic solution (09366-44: Nacalai Tesque). After 10 min, the PEG medium in the chamber was carefully replaced with PBS++ without PEG. Then, 4 *μ*l of 1.3 *μ*g/*μ*l eGFP-histone H1A or 1.5 *μ*g/*μ*l mCherry-histone H1A in Dulbecco’s phosphate-buffered saline without Ca^2+^ and Mg^2+^ (PBS--) was applied to the cuvette of a purpose-built electroporator and was directly embedded in the migrating keratocytes as described previously^45^. Next, the medium in the chamber was replaced with the culture medium. The keratocyte cultures were then left overnight to allow expression of eGFP- or mCherry-histone H1A.

### Staining of actin filaments in live keratocytes

Alexa Fluor 488 phalloidin (A12379: Life Technologies) in PBS++ or Alexa Fluor 546 phalloidin (A22283: Life Technologies) in PBS++ were introduced into the migrating keratocytes using the purpose-built electroporator^45^. SPY555 (CY-SC202: Spirochrome) was used in some experiments (Fig. 6B). It was dissolved in 50 *μ*l DMSO, and diluted 1:500 in culture medium. This SPY555 medium was then added to the chamber, to the bottom of which the cells had adhered, just after removal of the culture medium. The chamber was kept at 37 ºC. After 30 min, the SPY555 medium in the chamber was carefully replaced with the culture medium. The cells were then allowed to recover for about 5 min before observation.

### Staining of the membrane in live keratocytes

Alexa Fluor 488 conjugate of Con A (C11252: Thermo Fisher) was dissolved at 50 *μ*g/ml in culture medium. This Con A-Alexa medium was then added to the chamber, to the bottom of which the cells had adhered, immediately after removal of the culture medium. After 10 min, the Con A-Alexa medium in the chamber was carefully replaced with the culture medium. The cells were then allowed about 5 min to recover before observation.

### Blebbistatin treatment

Blebbistatin treatment was performed as previously described^7,25^. Briefly, (±)-Blebbistatin (13186; Cayman) was dissolved at 100 mM in DMSO and then diluted 2,000 times with culture medium to a final concentration of 50 *μ*M. This blebbistatin medium was then added to the chamber, to the bottom of which the cells had adhered, immediately after removal of the culture medium. After 30 min, the cells in the chamber were used for experiments without removal of the blebbistatin medium.

### Microscopy

The migrating keratocytes were observed using an inverted microscope (Ti; Nikon) equipped with a laser confocal scanner unit (CSU-X1; Yokogawa) and a high-speed z-axis scanner (NZ100CE, Prior) through a 100 × objective lens (CFI Apo TIRF 100 × H/1.49: Nikon). The fluorescence images were detected using an EM CCD camera (DU897; Andor).

About 40 slices of the optical sections were recorded at 0.5-*μ*m intervals to construct a 3D cell image. The time interval for recording each optical section was 56 msec. The consecutive images of each optical section were reconstructed into 3D images using FluoRender (SCI, Univ. of Utah). The image data were analyzed using Fiji (https://fiji.sc).

### Construction of mechanical model

A mechanical model (Fig. 3A and B) which mimics the keratocyte’s cell body containing stress fibers was made as follows. The main body was made from a type of PDMS (CY52-276A and B, Dow Corning Toray). A mixture of CY52-276A and B at a weight ratio of 7:10 was poured into a cylindrical polypropylene mold (14 mm in inner diameter and 30 mm long) and allowed to solidify at room temperature (25 °C) for two days. Before solidification, a 20-mm long coiled shape-memory alloy (SMA; BMX150, Toki Corp.) was extended to 30 mm and embedded in the surface layer of the body in the length direction. The Young’s modulus of the gel was measured using the method of Lo et al.^46^. It was estimated to be 6.3 kPa for a mixture ratio of 7:10. In response to the application of electric current to the expanded SMA, it contracts due to the self-generated heat as described elsewhere^47^. Electric current was applied to the coil using a purpose-built electric power circuit (Supplementary Fig. 1B).

For comparison with the mechanical model, another mobile with a frontal self-propelled wheel and a rear passive wheel was also made. The wheels were made from acrylic plastic. Their diameter and length were the same as the model. The power source of the self-propelled wheel is a motor with low-speed gear (70189, Tamiya).

### Traction force measurements

Traction forces of migrating keratocytes were measured as previously described^13,48^ with minor modifications. Briefly, CY52-276A and B were mixed at a weight ratio of 6:10. A 40 - 45-mg aliquot of the mixture was spread on a 22 × 22 mm coverslip (No. 0, Matsunami). After the mixture had solidified at room temperature (25 °C) for two days, the substrates were kept in a hermetically sealed case with a 50-ml aliquot of liquid silane (3-aminopropyl triethoxysilane, Sigma-Aldrich) at 70 °C for 1 h, to attach the silane to the surface of the substrate by vapor deposition. Fluorescent carboxylate-modified beads (0.1 *μ*m in diameter; F-8800, Life Technologies) were then attached to the surface of the substrate. The Young’s modulus was estimated to be 13.8 kPa for the mixture ratio of 6:10.

To measure the traction forces exerted by the mechanical model, a large elastic sheet of substrate was made. A mixture of CY52-276A and B at a weight ratio of 8:10 was poured into a dish (90 mm in inner diameter and 15 mm in height) and allowed to solidify at room temperature (25 °C) for two days. Another mixture of CY52-276A and B at a ratio of 8:10 containing zirconia balls (0.5 mm in diameter) was then poured on the top of the previous layer and allowed to solidify at room temperature (25 °C) for two days. The thicknesses of the bottom and top layers were 10 mm and 0.5 mm, respectively. The Young’s modulus was estimated to be 5.1 kPa for the mixture ratio of 8:10.

The traction forces exerted by keratocytes or the mechanical models were calculated from the displacements of the beads or the zirconia balls using Fiji and its two plug-ins, PIV and FTTC^49^. The regularization parameters were set at 3 × 10^−10^ and 3 × 10^−8^ respectively for the keratocyte and mechanical model traction force reconstructions.

### Detection of pushing against the substrate

A soft elastic substrate was made in the same manner as the traction force measurements of the mechanical model, except that the zirconia balls were embedded in a straight line. The mechanical model was placed on the line (Fig. 5I). The height of the space in which the mechanical model could move was limited to 14 mm, the same as the diameter of the gel, with an acrylic plate. The displacements of the zirconia balls caused by the movement of the model were recorded from directly behind the model.

A soft elastic substrate was made in the same manner as the traction force measurements of keratocytes, except that CY52-276A and B were mixed at a weight ratio of 10:7. The Young’s modulus was estimated to be 0.1 kPa for the mixture ratio of 10:7. Keratocytes were dispersed on the substrate. The height of the space where the cells can move was limited to their height by placing a 1% agarose (01163-76; Nacalai Tesque) gel sheet (5 mm × 5 mm × 1 mm) on them (Fig. 6A). About 30 slices of the optical sections of the fluorescent bead images were recorded at 0.3-*μ*m intervals to construct a 3D image. The height of the beads was analyzed using a self-made Python script.

In both the mechanical model and keratocytes, in order to estimate the pushing force using the method of Lo and colleagues^46^, we assumed that two spherical protrusions push the substrate. The pushing force *F* was calculated as *F = 4d*^*3/2*^*r*^*1/2*^*Y/3(1-p*^*2*^), where *Y* is the Young’s modulus of the substrate, *d* is the indentation, *r* is the radius of the spherical protrusion, and *p* is the Poisson ratio (assumed to be 0.5^50^). For keratocytes, focal adhesions were considered as the spherical protrusions, and their cross-sectional area was set to be 24.3 *μ*m^2^ from our previous study^18^.

## Acknowledgements

We thank K. Katoh (AIST, Japan) for kind gift of eGFP- and mCherry- histone H1A expression plasmids. Y.I. was supported by Innovative Science and Technology Initiative for Security, ATLA, Japan and MEXT Kakenhi Grants Nos. 22H05683, 21K19228 and 19H04935. Y.I and T.S. were supported by MEXT Kakenhi Grants No. 20H03227. C.O. was supported by MEXT Kakenhi Grants No. 21K15055. Y.N. was supported by MEXT Kakenhi Grants No. JP21H05308. We are grateful for their support.

## Author contributions

C.O. and Y.I. designed the research and wrote the manuscript. C.O. performed the experiments using the keratocytes. C.O., S.A., R.Z. and Y.I. constructed the mechanical model and performed the experiments using it. Y.N. developed the Python script for image processing. C.O., Y.N. and T.S. analyzed the experimental data. All the authors edited and approved the manuscript.

## Data availability

All raw data generated in this study are available on Zenodo, at https://doi.org/10.5281/zenodo.7502798.

## Notes

https://doi.org/10.5281/zenodo.7502798

### Competing Interest Statement

The authors have declared no competing interest.

