## Supplementary Information for "Linear contraction of stress fibers generates cell body rotation"

**This PDF file includes:**

Supplementary Figures 1 to 4

Legends for Supplementary Videos 1 to 12

**Other supplementary materials include the following:**

Supplementary Videos 1 to 12

#### Supplementary Figures

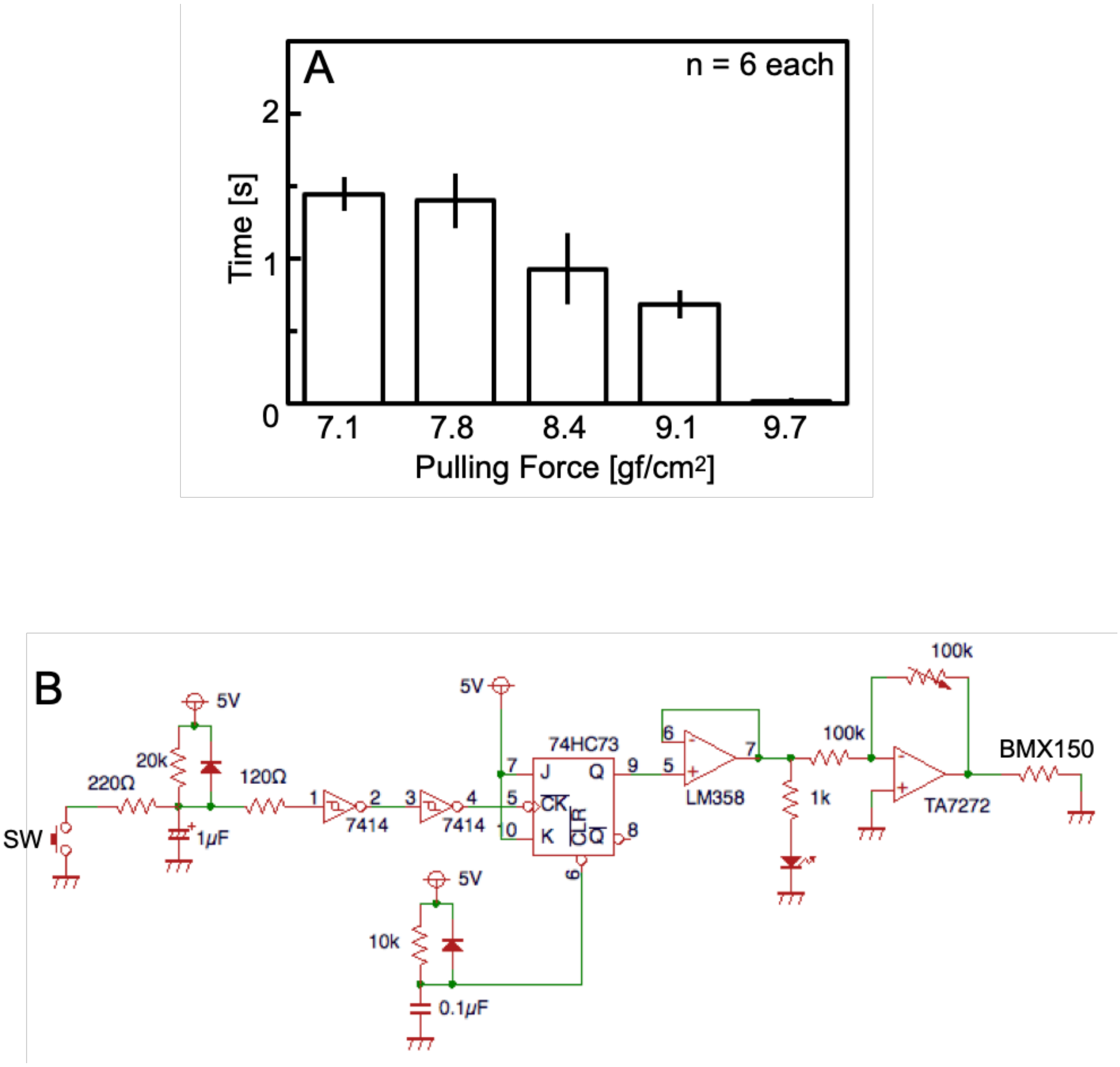


**Supplementary Figure 1.** Supplemental details of the mechanical model. (A) Time required for a mechanical model adhered to a substrate to detach from it for each pulling force. (B) Purpose-built electric power circuit for applying current to the coil (BMX150) by pushing the switch (SW). Error bars in A represent SEM. The *P* values were calculated using Student's *t* test.

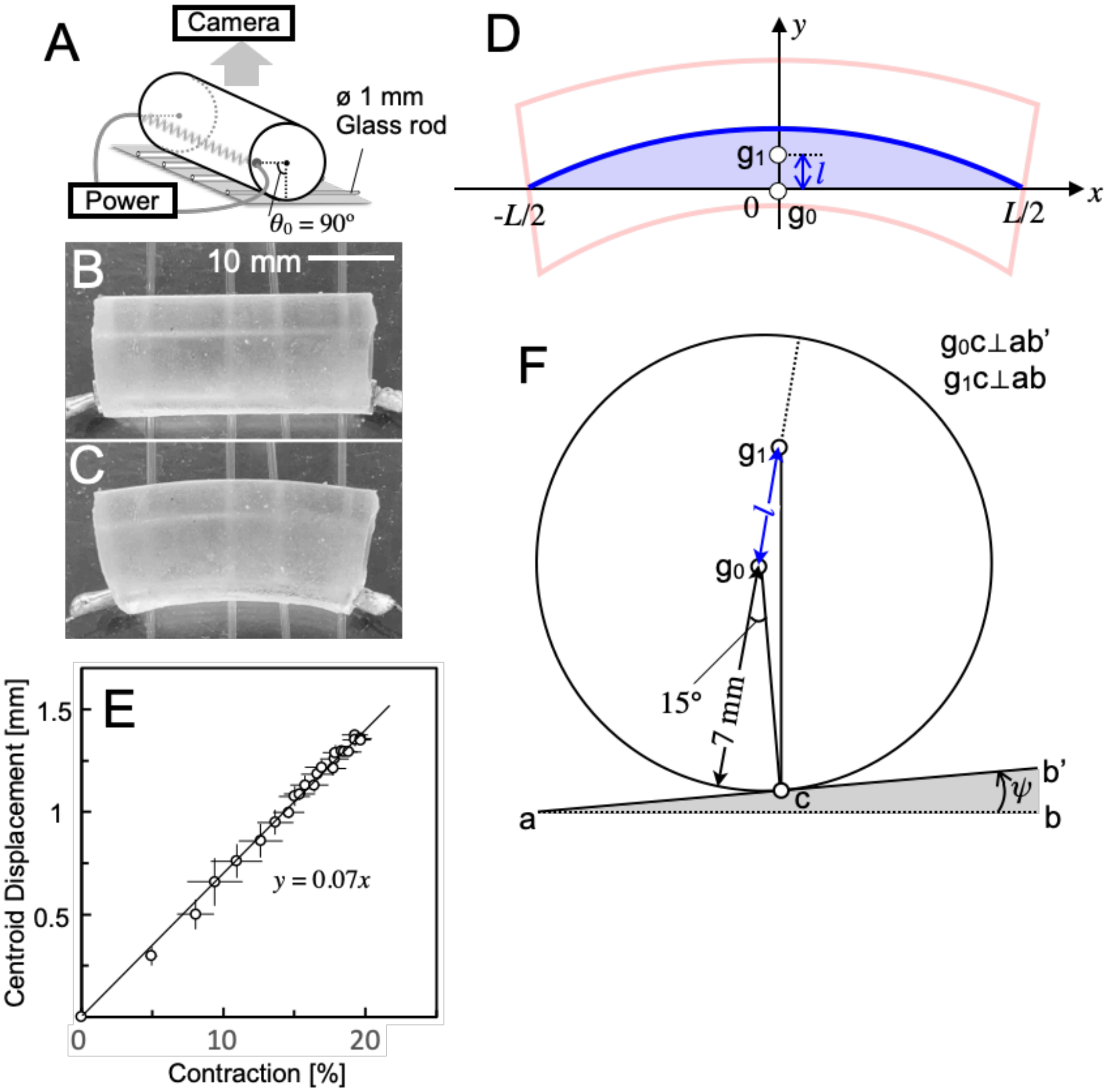


**Supplementary Figure 2.** Estimation of the displacement of the centroid of the gel due to the contraction of the coil and the tilt angle of the substrate to cancel it. (A) Schematic illustration of the model deformation recording. (B and C) Images of the model before (B) and after (C) application of the electric current to the coil. (D) Estimation of the displacement of the centroid ($l$). (E) Estimation of tilt angle ($\psi$) of the substrate. When the substrate is tilted $\psi$ degrees, the gravitational pull on the new centroid position should align with the grounding point of the gel.

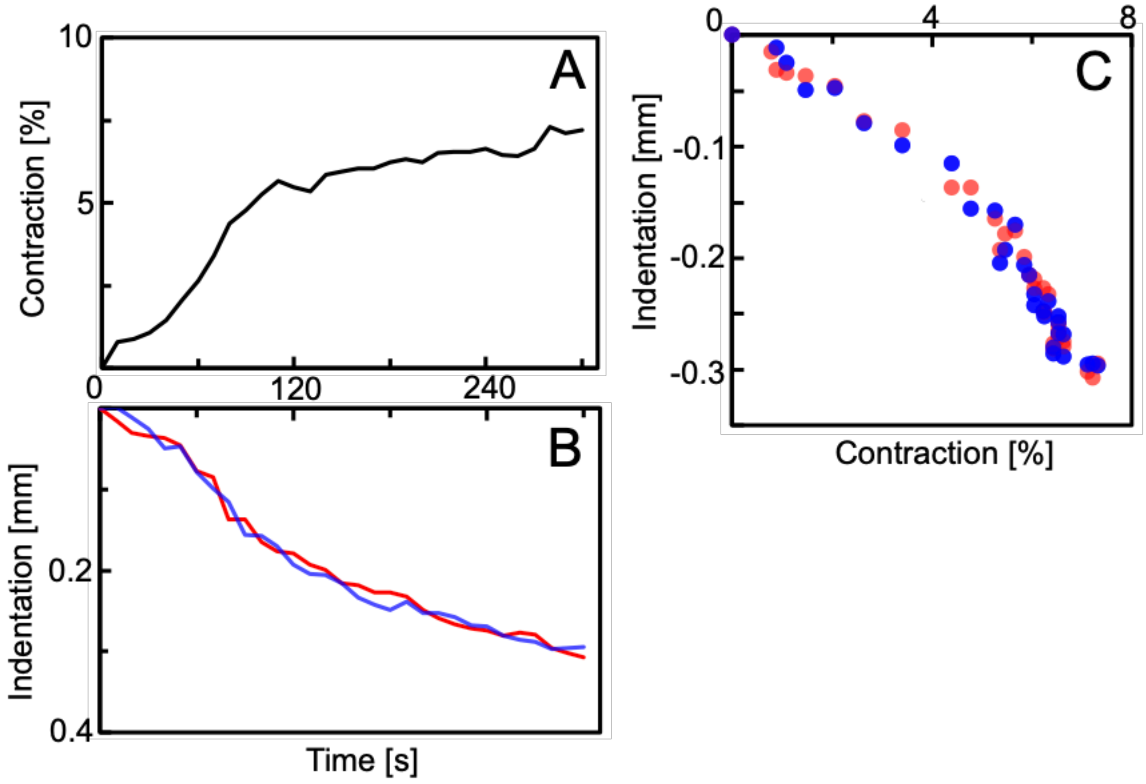


**Supplementary Figure 3.** Indentation of soft substrate due to the motion of the mechanical model. (A and B) Time courses of the coil contraction of the model (A) and indentation of the substrate just below the left (red) and right (blue) edges of the model (B). (C) Contraction-indentation curve made from (A) and (B).

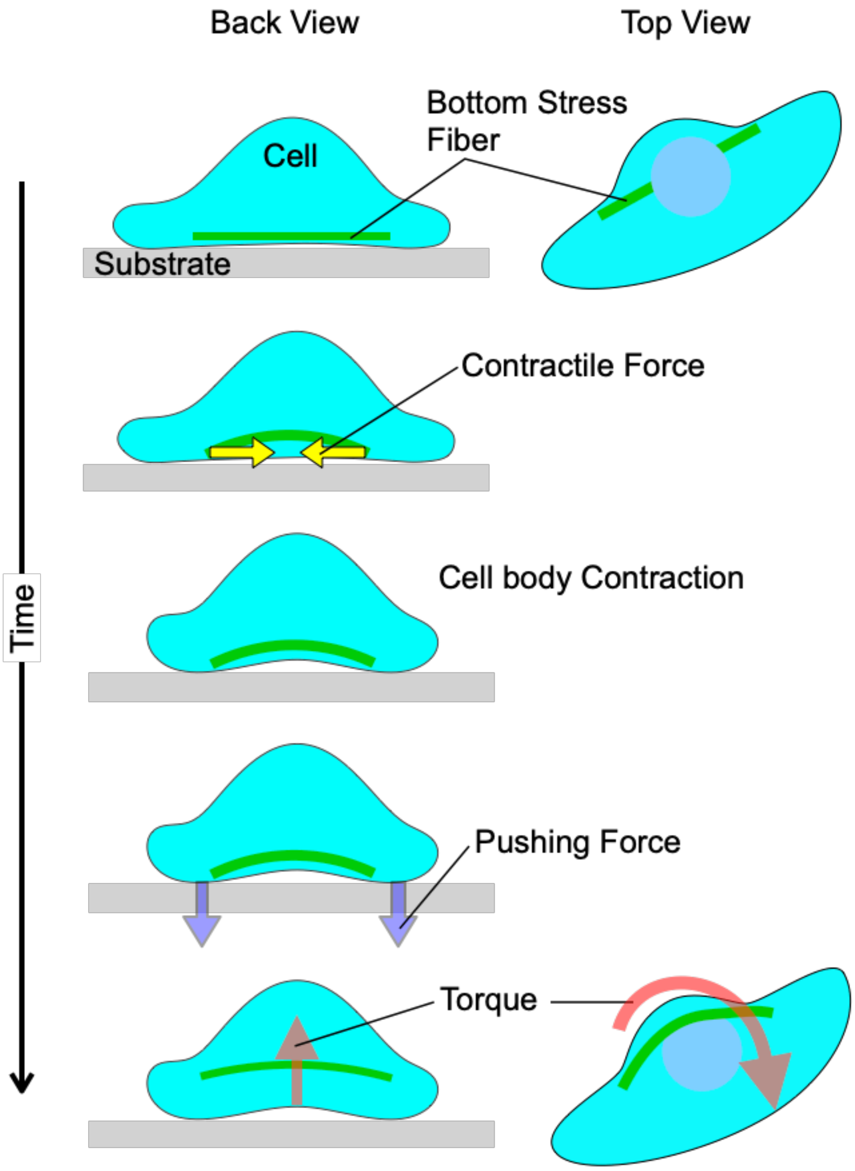


]**Supplementary Figure 4.** Conclusion of this study: how linear contraction of stress fibers is converted into rotation. Contraction of stress fibers deforms the cell body, and the deformed cell body pushes against the substrate. The pushing force generates the torque for rotation.

### Legends for Supplementary Videos

**Supplementary Video 1.** A migrating keratocyte. The video depicts the same keratocyte as that shown in Fig. 1A and is shown 45 times faster than real time.

**Supplementary Video 2.** Sequential 3D fluorescence images of GFP-histone H1A in a migrating keratocyte. The video depicts the same cell as that shown in Fig. 1C - F, and is shown 75 times faster than real time.

**Supplementary Video 3.** Sequential images of the nucleus and the actin cytoskeleton in a migrating keratocyte. Top: 3D images. Bottom: perpendicular optical sections. Fluorescence of GFP-histone (magenta) and Alexa Fluor 546 phalloidin (Green). White arrow: a stress fiber. Yellow arrow: the protrusion of a slightly distorted nucleus. The video depicts the same cell as that shown in Fig. 1H - J, and is shown 80 times faster than real time.

**Supplementary Video 4.** Sequential images of the cell membrane of a migrating keratocyte. Fluorescence of Con A-Alexa at the top (Top) and bottom (Bottom) optical sections of a migrating keratocyte. The identical Con A spots are indicated by same-color arrowheads on each panel. The video depicts the same cell as that shown in Fig. 2A - D, and is shown 25 times faster than real time.

**Supplementary Video 5.** Sequential images of the actin cytoskeleton and cell membrane at the bottom optical section of a migrating keratocyte. Fluorescence of Alexa Fluor 546 phalloidin (F-actin), Con A-Alexa (Membrane) and the two merged (Merged). Yellow arrows and circles indicate a stress fiber and a single Con A spot, respectively. The video depicts the same cell as that shown in Fig. 2F - K, and is shown 30 times faster than real time.

**Supplementary Video 6.** Movement of a mechanical model of a keratocyte cell body. Left: side view. Right: rear view. The yellow arrowheads indicate the positions of the coil in the first frames. The video depicts the same mechanical model as that shown in Fig. 3D and E, and is shown 150 times faster than real time.

**Supplementary Video 7.** Traction forces exerted by the mechanical model. The video depicts the same model as that shown in Fig. 4A and is shown 300 times faster than real time.

**Supplementary Video 8.** Traction forces exerted by a migrating keratocyte. Force images (Force) and DIC images (DIC). The video depicts the same keratocyte as that shown in Fig. 4B and is shown 60 times faster than real time.

**Supplementary Video 9.** Traction forces exerted by a mechanical model of a vehicle containing a driving front wheel and a non-driving rear wheel. The video depicts the same model as that shown in Fig. 4D and E, and is shown 2.5 times faster than real time.

**Supplementary Video 10.** Deformation of a soft substrate (Young's modulus: 5.1 kPa) with zirconia balls (0.5 mm in diameter) embedded in its surface by the contraction of the coil in the mechanical model. The red and blue circles are the positions of the balls on each side of the model in the first frame. The video depicts the same mechanical model as that shown in Fig. 5J - N, and is shown 150 times faster than real time.

**Supplementary Video 11.** Sequential perpendicular optical sections of the actin cytoskeleton in a migrating keratocyte. On a soft substrate (Young's modulus: 0.1 kPa) (top, SPY555) and on a coverslip (bottom, Alexa-phalloidin). The optical section is parallel to the migration direction at the center of the cell. The yellow arrows indicate single stress fibers. The video depicts the same cell as that shown in Fig. 6B, and is shown 40 times faster than real time.

**Supplementary Video 12.** Local indentation of a soft substrate (Young's modulus: 0.1 kPa) beneath the left and right ends of the cell body of a migrating keratocyte. The indentation was detected using fluorescent beads (0.1 $\mu$m in diameter) embedded in the surface of the substrate. DIC images (DIC) and fluorescent bead images (Beads). Three beads passing under the left, middle, and right of the cell are indicated by the three colored squares. The video depicts the same cell as that shown in Fig. 6C, and is shown 50 times faster than real time.
